# Improving Interdisciplinary Teaching through a Complexity Lens

**DOI:** 10.1101/2022.05.26.493642

**Authors:** Sarah Neitzel, Yuhao Zhao, Carrie Diaz Eaton

## Abstract

In this article, we discuss the use of bipartite network analysis to understand and improve interdisciplinary teaching practice. We theorize mathematics and biology as part of a coevolving mutualistic ecosystem. As part of an interdisciplinary teaching initiative, we inventoried mathematics topics appearing in the marine biology classroom and their associated marine context. We then apply techniques of mutualistic bipartite networks analysis to this system to understand the use of mathematical concepts in a marine biology classroom. By analyzing the frequency and distribution of mathematics topics, we see that a variety of mathematical concepts are used across the course with most appearing only a few times. While this is an inherent trait of mutualistic coevolutionary networks, it can create a logistical challenge to supporting mathematics in the marine biology classroom. We find that marine biology topics containing the most mathematics are either close to the instructor’s research area or were introduced through externally developed educational resources. Finally, we analyze groups of topics that appear connected to each other more frequently to inform both interdisciplinary education development as well as disciplinary support. We also suggest ways to use network metrics to track interdisciplinary connections over time, helping us understand the impact of interventions on interdisciplinary teaching practice.

## INTRODUCTION

### Disciplines as a coevolving and mutualistic complex adaptive system

As new interdisciplinary fields emerge, interdisciplinary education has become vital. Students are expected not just to know concepts in their own field, but to be able to apply them in novel situations outside one’s discipline. For biology education, this expectation is clearly delineated in the *Vision and Change* report’s description of core competencies and disciplinary practice (AAAS, 2011). This report defines expectations for students to harness the “interdisciplinary nature of science” and “collaborate and communicate with other disciplines’’ (p. 15), as well as specific expectations for developing skills in quantitative reasoning, modeling, and simulation (AAAS, 2011). A driver of these educational policy conversations was the emergence of mathematical and computational biology as a recognized and highly active subdiscipline at the turn of the century (Diaz Eaton et al., 2020).

*Vision and Change* as an educational policy report is an example of textual artifact that reflects the response of biology education to an increasingly mathematical and computational research agenda. These educational calls are reiterated in other biology education policy reports such as *Bio2010*, by the National Research Council (NRC, 2003), which outlines mathematical curriculum. But also, this response is mirrored in mathematics education policy reports, for example Guidelines for the Assessment and Instruction of Mathematical Modeling Education (Garfunkel & Montgomery, 2019), which was a response to interest in creating more opportunities for studying mathematical modeling throughout the mathematics curriculum.

In this paper, we theorize mathematical biology education as an emergent field resulting from the interactions over time of two mutualistic coevolving disciplines, biology, and mathematics. Specifically, we theorize these interactions as mutually beneficial. Biology research informs interesting mathematical questions - for example, how can pollinator networks evolve with overwhelming specialism, but remain evolutionarily resilient (Bascompte & Sheffer, 2023)? Likewise, mathematical research informs interesting biology questions - for example, how does speciation happen within populations without geographic isolation (Bolnick & Fitzpatrick, 2007)? In the context of education at the interface of mathematics and biology as instructors work to infuse an interdisciplinary perspective across courses, we may see mathematics instructors using biological observations to motivate mathematical investigations and/or interpret the findings. In a biology classroom, we may see instructors using mathematical ideas and tools to explain or explore observed biological phenomena. Authentic integration of content across interdisciplinary boundaries increases students’ perceived utility value of the discipline and may contribute to more positive affect and performance (Aikens et al., 2021).

There have been a number of course reforms from infusing biology examples into calculus and to modeling in introductory biology as well as new biology or mathematical biology courses at the interface (Diaz Eaton & Callender Highlander, 2017; Robeva et al., 2022; Weisstein, 2011; Acevedo, 2020). There are also a number of funding initiatives and communities of support which support mathematics and biology instructors to meet the demand for a more integrated education (Diaz Eaton et al., 2020; Akman et al., 2020). Nonetheless, it remains a challenge to help instructors who have not been cross trained in both disciplines to teach in interdisciplinary ways, for a variety of reasons including differences in epistemology and language (Redish & Kuo, 2015; Diaz Eaton et al., 2019).

Henderson, Beach, and Finkelstein (2011) reviewed STEM education reform strategies with particular attention on teaching practice intervention. These interventions typically operate on two axes: intervening at the environment or the individual and using prescribed or emergent interventions. However, most of the current work on the assessment of interdisciplinary thinking is focused on undergraduate students (Mansilla et al., 2006; Mansilla & Draising, 2007; Spelt et al., 2009). While this is an important outcome to assess, it does not tell us explicitly about how teachers might come to their interdisciplinary teaching practice, nor how the curriculum is evolving to become more interdisciplinary. Viewing the curriculum, or rather artifacts of the curriculum such as lecture or texts, as snapshots of a broader co-evolutionary process between ideas across disciplines sampled and enacted by instructors may give us new insight into interdisciplinary education.

We use the context of a marine biology course, and the lectures presented, to examine a network of mathematical and biological ideas in the context of a broader interdisciplinary professional development. More specifically, our sample includes a qualitative examination of the lectures of an introductory marine biology course for calculus “ideas.” These calculus and marine biology “ideas,” form the nodes of the lexical network and co-occurrence between a calculus and a marine biology idea is a tie or link. Our research questions are:

- Is there structural evidence from the lectures that the network of mathematical and biological concepts may have arisen from a broader mutualistic complex adaptive system between mathematics and biology?
- How many calculus concepts arise in an introductory marine biology course, and what kinds of topics are important?
- With a network science and complex adaptive systems lens on the analysis of this concept network, what new insights can be derived about interdisciplinary education, both for our specific marine-math network, and more broadly?

In education research, network analysis tends to focus on relationships between individuals, both teachers and students, or the affiliations of individuals (Grunspan et al., 2014; Dou et al., 2016, and Buchenroth-Martin et al., 2017; Andrews et al., 2016). An affiliation network approach has also been used to assess interdisciplinary research, typically through examining citation networks (Palmer, 2013; Bishop et al., 2014). Possibly analogous to understanding interdisciplinary teaching professional development, these results have been used to understand how synthesis centers help change research practice to become more interdisciplinary (Baron et al., 2017).

While transcription and qualitative coding of course content and lectures is a common approach in education research (e.g., Choe et al., 2019), computational text analysis versions are becoming more common in STEM education (e.g., Moreno et al., 2019). However, network science approaches to conceptualizing and analyzing lexical database are still limited and primarily pioneered by scholars in digital humanities (e.g., Koponen, 2021). Network approaches which emphasize exploring the structure and resilience of networks as physical complex adaptive systems are common in research on biological evolution (Proulx et al., 2005; Ings et al., 2009) and in research on the evolution of language networks (Solé, 2010; Greenhill, 2010; List, 2013).

There is also a significant amount of scholarship in “knowledge systems” which builds theories of learning on individual’s organization of concepts and evolution and transitions from weak organization to more sophisticated levels of organization (diSessa & Sherin, 1998; Sabella & Reddish, 2007). Some of these studies include an explicit network scheme, sometimes referred to as a semantic net, which relate concept associations to student learning (Marshall, 1995, p.192; Eremin, 2014).

Closest to the methodology in our study is Koponen (2021), which uses a complex adaptive systems frame and network approaches to examine disciplinary ideas in the History of Science. In this context, a unimodal network analysis was employed to examine topics and their relationships within that subject. Kopenen uses textual documents and employs computational techniques to form the lexical database of topics, whereas we rely on qualitative coding techniques of verbal lectures. Also, because we are focused primarily on the relationships between two disciplines, marine biology and calculus, we use bipartite network analysis.

Because we assume mutualism between interactions between mathematics and biology, we approach the network questions similarly to other systems which analyze coevolving bipartite mutualistic networks in biology (Bascompte and Jordano, 2007; Ings et al., 2009; Vázquez et al., 2009).

## METHODS

From 2016 - 2018, Unity College, a small environmental liberal arts college, was part of a ten-college networked improvement community effort, SUMMIT-P, to revise foundational mathematics courses. The idea was to partner with, not for, partner disciplines to improve mathematics learning outcomes for all students (Beisiegel & Doree, 2020). Each college within the SUMMIT-P consortium has at least one co-PI from mathematics, a target foundational mathematics course or courses, and at least one co-PI from a partner discipline. Unity College’s project was to revise the Calculus I and II sequence to meet the needs of the Marine Biology major. Calculus I and II had already adopted scientific writing which utilized the same scientific writing rubric as the Biology and Marine Biology classes (Diaz Eaton & Wade, 2014) and had revised the content to meet the needs of the two programs which required it: Wildlife Biology and Earth and Environmental Science (Diaz Eaton & Callender Highlander, 2017). In Fall 2016, the Marine Biology program began requiring Calculus I as part of its major, thus its interest in a partnership between the Calculus courses and the Marine Biology program.

As part of this grant, the Marine Biology program director sat in on Calculus I, and the next semester, the Calculus instructor (co-author Diaz Eaton) sat in on Introduction to Marine Biology, a sophomore course for Marine Biology majors and elective for other majors.

Introduction to Marine Biology, hereafter referred to as just “Marine Biology,” met for 50 minutes on Wednesdays and Fridays for lecture and then for an additional 4-hour lab on Friday afternoons. Diaz Eaton has graduate training in both mathematics, ecology, and evolution. At the time of data analysis, Neitzel was an undergraduate in marine biology with a minor in Applied Mathematics and Statistics and served as the undergraduate student TA for Calculus and a research assistant on the project. Yuhao Zhao, an undergraduate mathematics and economics major, joined the project during the later stages of network analysis.

Lecture data was collected by Diaz Eaton starting on the second lecture (the first was a syllabus review) and was not collected during lab periods. Instead, during the lab periods, Diaz Eaton acted as an assistant in the lab classroom. During the lecture portions, when a math concept was recognized as being used in relation to the marine topic at hand (explicitly or implicitly), it was recorded, along with the context in which it occurred, as illustrated in Table 1. As a pilot project, we were looking for emergent themes and keywords across a broad mathematics spectrum.

**Table 1.**
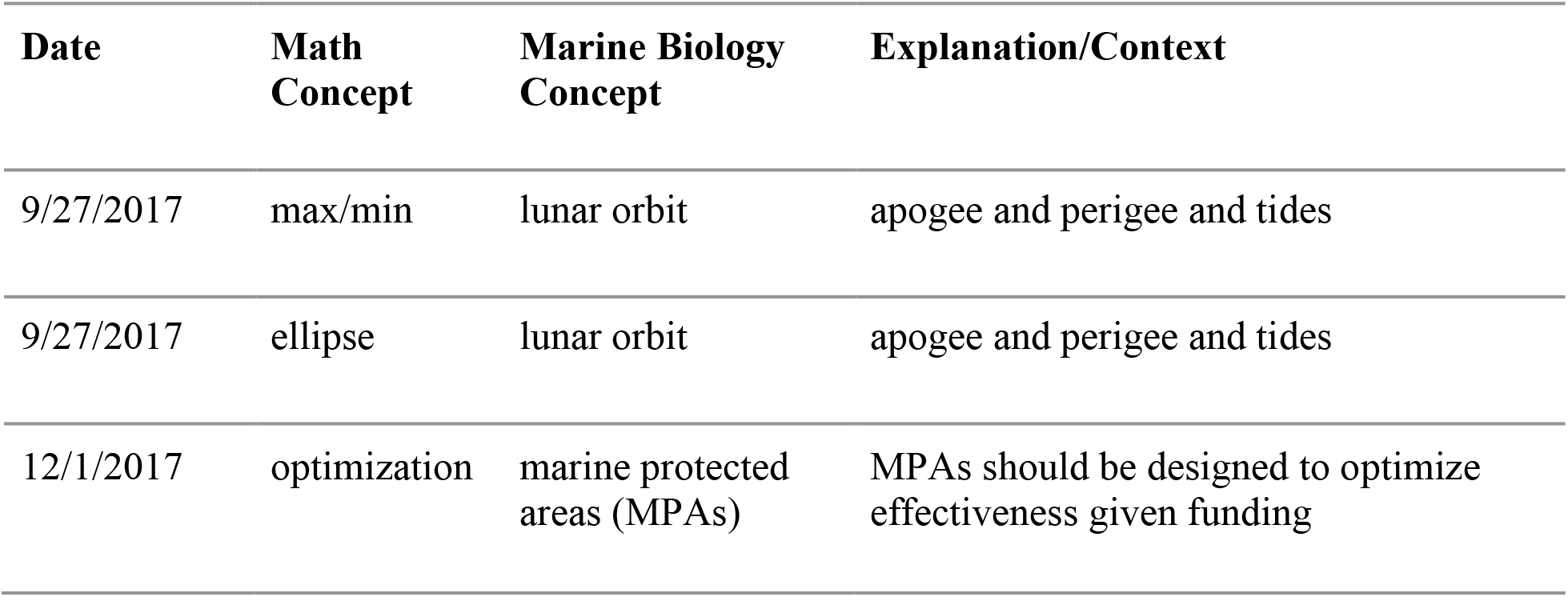
Simplified excerpt of data table of math and marine concept interactions.

Therefore, we did not establish an *a priori* ontology, but rather allowed it to emerge as a result of the observations. For boundaries around which mathematical terms were recorded, we limited ourselves to the concepts present in the Unity College version of Calculus I and II (Diaz Eaton & Highlander, 2017), as well as their prerequisite algebra courses. It should be noted that the algebra through calculus curriculum at Unity has a heavy emphasis on modeling. Calculus I includes multivariable visualization and differentiation, so terms such as “scale” or “3D,” fall within our “calculus” tent. We excluded mathematical terms that were statistical (e.g., hypothesis), but it should be noted that statistical language and ideas were frequently used in the lab component of the marine biology course. Due to our broad definition of “calculus” terms for this context, we refer to these terms more broadly as “math.” Recordings of “math” concepts were often due to explicit use of the terminology in class, or use of similar phrases. For example, the word “optimize” was used in context of MPAs, but the term “optimal” was used in the lecture on evolution under constrained salinity systems. Both were ultimately recorded under the umbrella concept “optimization.”

As part of data manual data cleaning and thematic analysis, Netizel, in consultation with Diaz Eaton, grouped topics if they shared word stems (e.g., graph and graphing), were subcategories of a broader topic (e.g., conservative systems were grouped under systems), or were otherwise closely related conceptually (e.g., min/max, optimal, and optimization). The result is a list of math and marine biology “concept” co-occurrences which form an edgelist for the bipartite network (edgelist data is in the CSV file in Supplementary Materials). Observation data taken within a single Marine Biology course is akin to plant-pollinator network observation data in which particular plant areas are observed for pollinators. The math concepts list should be considered complete, whereas the marine concepts list should be considered incomplete, as only marine concepts with clear math connections were listed. This can skew data signals from the full underlying coevolutionary network (Gibson et al., 2011). For example, the lecture on the physics of tidal systems was only one day but is highly overrepresented in our dataset because of the number of math references within that one lecture period.

For the bipartite network analysis, we utilize the ‘bipartite’ package in R, which is commonly used in analysis of mutualistic complex adaptive systems.(Dormann et al., 2008; Dormann, 2020; Saavedra et al., 2009). To distinguish the two types of nodes in the bipartite system, ‘bipartite’ utilizes the terminology ‘lower’ and ‘higher’. We define ‘lower’ as mathematics and ‘higher’ as marine biology. We utilized a variety of network analysis approaches common to ecological bipartite analysis including network visualization, degree distribution, and clustering analysis of the interaction matrix to gain insights. The CSV data file and the RMD file are uploaded to GitHub(https://github.com/mathprofcarrie/marine-math-network).

## RESULTS AND DISCUSSION

### Characterizing which math is in the marine biology classroom

Figure 1 is a plot of the generated bipartite network data. When we visualize these connections, we notice immediately that there is not a clean one-to-one relationship between math topics and marine biology topics. The thickness of an edge indicates the strength of the connection (number of times that particular combination was mentioned). However, in the case study, most concepts were connected only once or twice. This is a common feature of network heterogeneity - few nodes have many connections, i.e., “generalists”, and many nodes have few connections, i.e., “specialists” (Vázquez et al., 2009).

**Figure 1.**
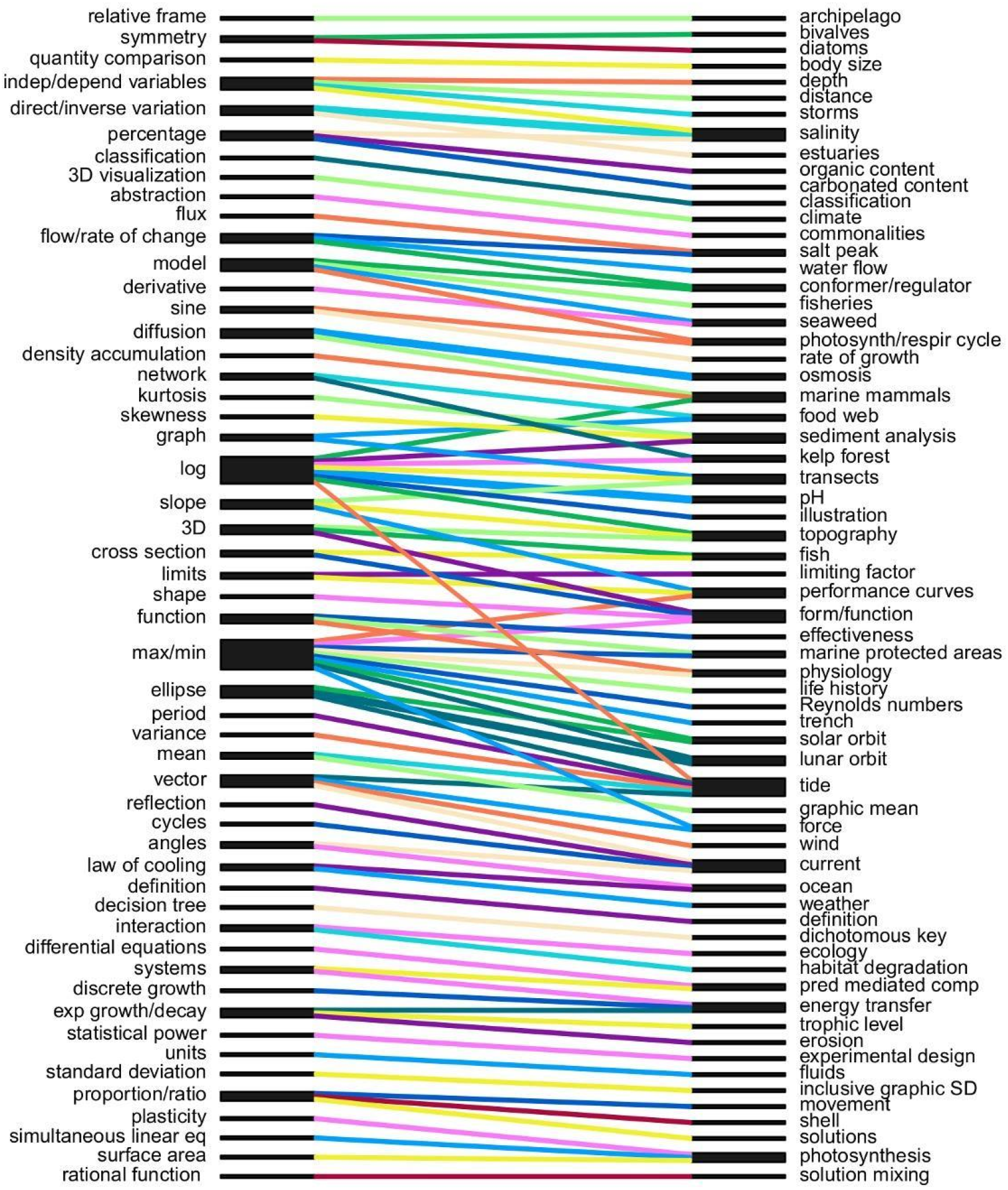
True web of math-marine concept network produced via ‘plotweb’. Math concepts (the ‘lower’ level) are on the right and marine concepts (the ‘higher’ level’) are on the left, utilizing ‘cca’ to group highly connected concepts together towards the center of the plot, which acts to minimize edge crossing. The black bars are nodes and the colored lines indicate co-occurring math and marine topics. The size of both indicates node or edge size respectively, with only node size varying dramatically in this web.

We can infer a few of the most common topics simply from the network graph. When we examine which mathematics topics appear most frequently, there are two topic areas that stand out. The grouping of themes related to maximum, minimum and optimization related to 10 different marine biology topics. The theme related to logarithms and scaling relate to 9 different marine biology concepts. On the other side of the web, you can see that tides are the “mathiest” marine concept. In the tail of Figure 1, there are four topics that are associated with the most mathematics: tides (6 connections), currents, salinity, and form and function (connections to 4 math topics each).

This study is a glimpse into an Introduction to Marine Biology classroom in which optimization and scale are stressed as key mathematical concepts. In this classroom, the instructor teaches concepts of tides, currents, salinity, and form and function in ways that are connected to many math topics. A good question to ask is whether these observations are inherent characteristics of introductory marine biology courses, or whether our observations signal something more important about the instructor and context. After re-examining lecture notes for these days, we find two interesting patterns.

Multiple mathematical references (e.g., max/min, mathematical function, dependent/independent variables, optimization) were made related to how organisms evolve in marine environments and the physiological adaptations needed to compensate for differing levels of environmental factors such as salinity, pressure. The instructor’s research area was in systematics and evolution, with an emphasis on marine invertebrate physiology. Recall, the instructor sat in on Calculus I in the previous semester, which included themes such as maximum, minimum, optimization, exponential growth, and log transformations (Eaton & Highlander, 2017). It may be that mathematics professional development took hold first in classroom topics close to the instructor’s research training.

The abundance of mathematical references in the lecture on tides and currents may be intrinsic to the discussion of physical oceanography. However, the level of explicit mathematical activity was highly influenced by the instructor’s choice to use an educational YouTube video produced by the National Oceanic and Atmospheric Administration to explain the tides. This video comprised a significant portion of the lecture period and included details about the physics.

Therefore, high quality third-party instructional materials may be a mechanism by which instructors are introducing quantitative concepts into the classroom.

These two strategies directly map onto the individual strategies proposed by Henderson, Beach, and Finkelstein (2011). The first strategy is emergent individual change, fostered by reflection and supporting new ways of thinking, which was a particular goal of the grant. The second is characterized as the use of a prescribed intervention for individuals.

## Distribution of Connections

Another way of conceptualizing the structure of connections is using a connectivity distribution (Figure 2). In connectivity distributions, “specialist” topics are indicated by the high bars on the left, but a long tail on the distribution, indicating that there are some “generalist” topics occurring multiple times. Both Figures 1 and 2 illustrate that, while there are some generalist topics that can be woven and reinforced into the math curriculum, there are also a number of specialist math topics which become important throughout the course.

**Figure 2.**
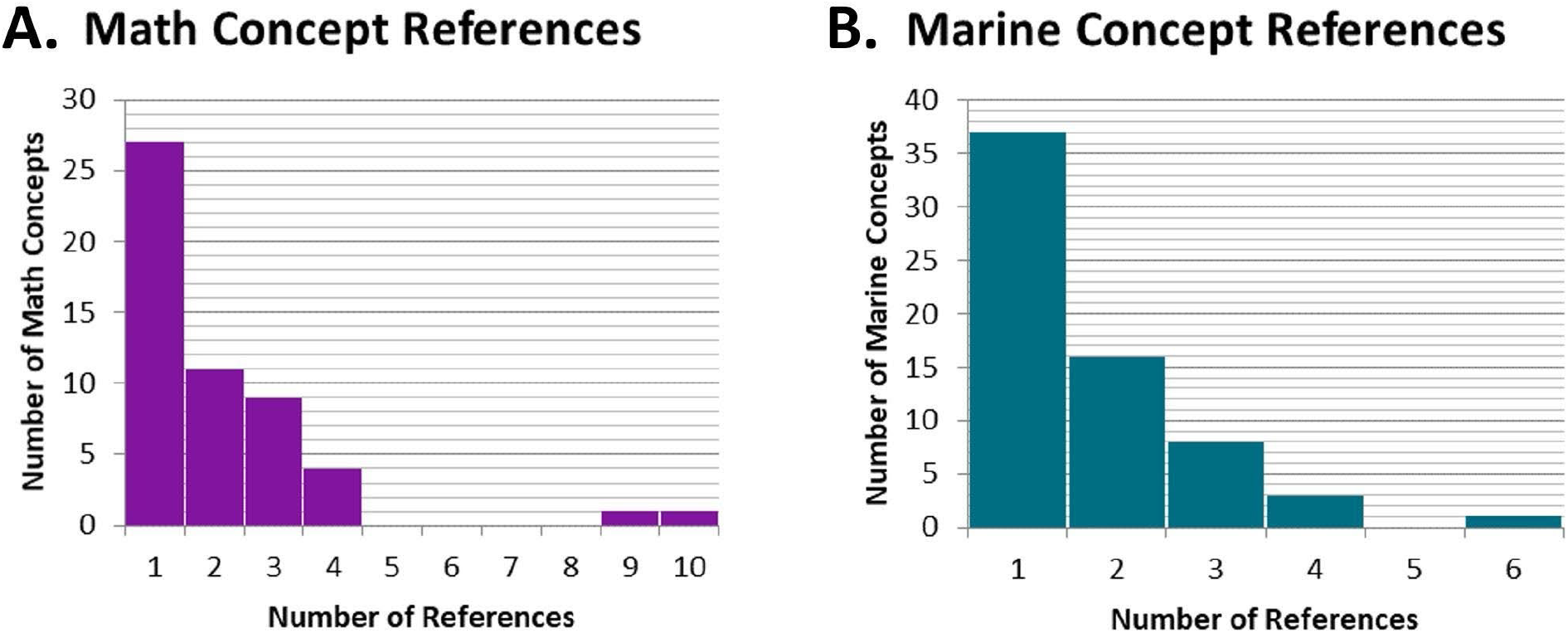
Histograms of the number of (A) math and (B) marine concepts referenced a given number of times during the Marine Biology course. In both cases, the majority of concepts were only referenced once (27 for math and 37 for marine) and frequency tapers off with only a select few concepts being referenced 4 or more times.

To gain insight from a student-centered perspective, we graphed the general flow of new and total math and marine concepts over the course of a semester (Figure 3). This shows that new math concepts and their related marine concepts generally spike at least once between exam intervals, then taper. The total number of concepts show a similar trend but with much more variation.

**Figure 3.**
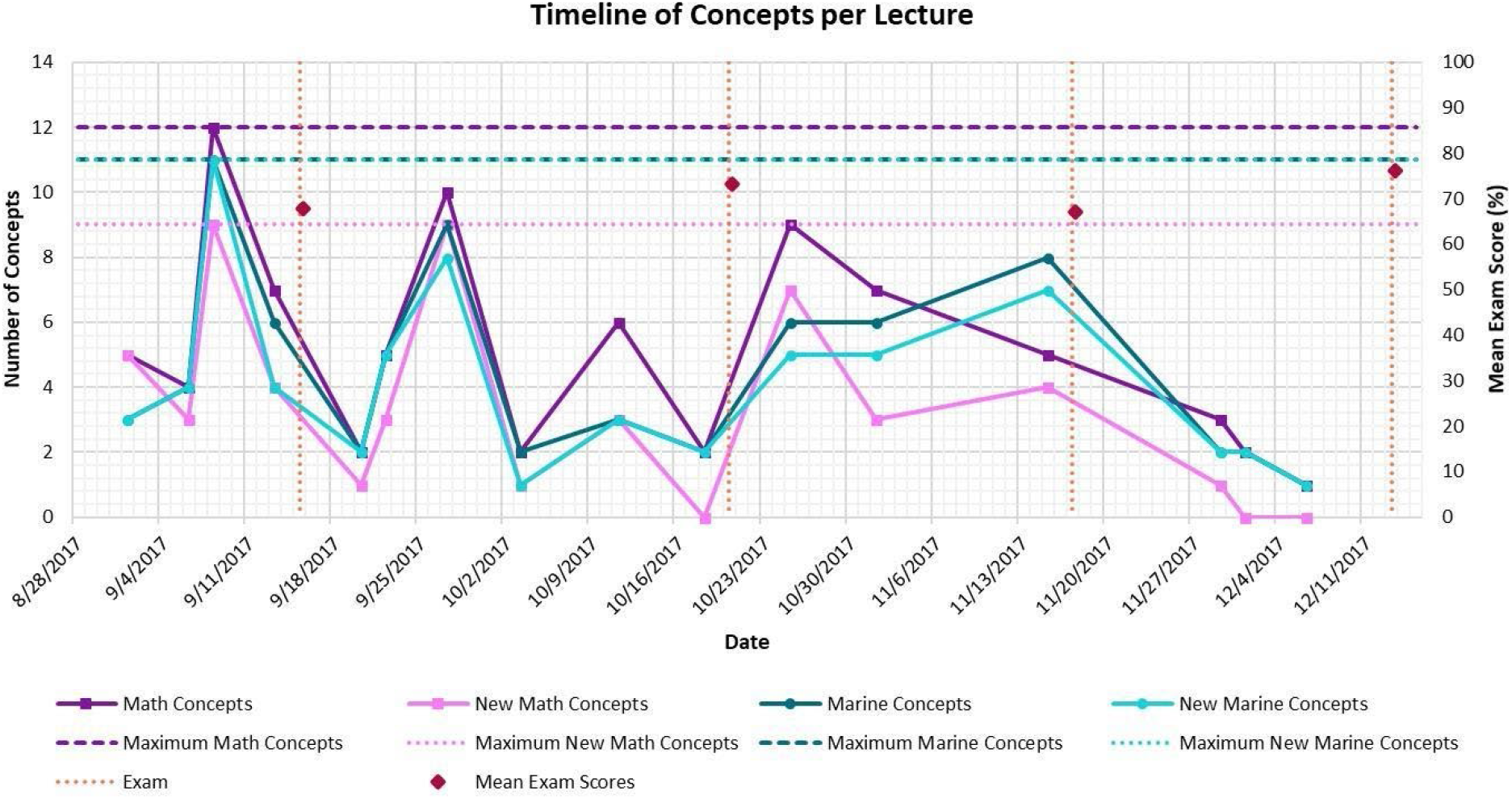
Timeline of Fall 2017 Marine Biology course with number of marine (teal/blue) and math (purple/pink) concepts covered per lecture along with how many were new vs. previously covered, maximum values, and exam dates (orange) and mean scores (red, as percentages).

We present this data as a cautionary tale for biology instructors who are wondering why their students have so much trouble with the mathematics in their classroom. In addition to issues of transference of mathematics out of the math classroom and into the marine biology classroom, we are expecting the transference of a wide array of mathematical concepts. In one semester of marine biology classroom lectures, we recorded 27 separate concepts of mathematics infused across 37 marine biology topics. Many of the topic areas were identified outside of the typical bounds for any one course or typical prerequisites. For example, vectors may be a topic relegated in some post-secondary curricula to be in Calculus III or perhaps in Physics I and may or may not be part of program requirements or class prerequisites.

### Supporting connections

By visualizing the interaction matrix between node types, we can better understand how topics support each other. In ecological networks, nested viswebs are used to examine nestedness (Bascompte & Jordano, 2007). In a highly nested network, all color in Figure 4A would be concentrated in the upper left corner and all or almost all ‘specialist’ concepts would only interact with ‘generalists’. If the web were highly nested, there would be a small number of ‘generalist’ concepts that could be easily focused on to increase interdisciplinary understanding. Our case study data were poorly nested: the interacting topics are highly dispersed, meaning that many specialists mathematical topics are scattered throughout many marine topics, a point also made above.

**Figure 4.**
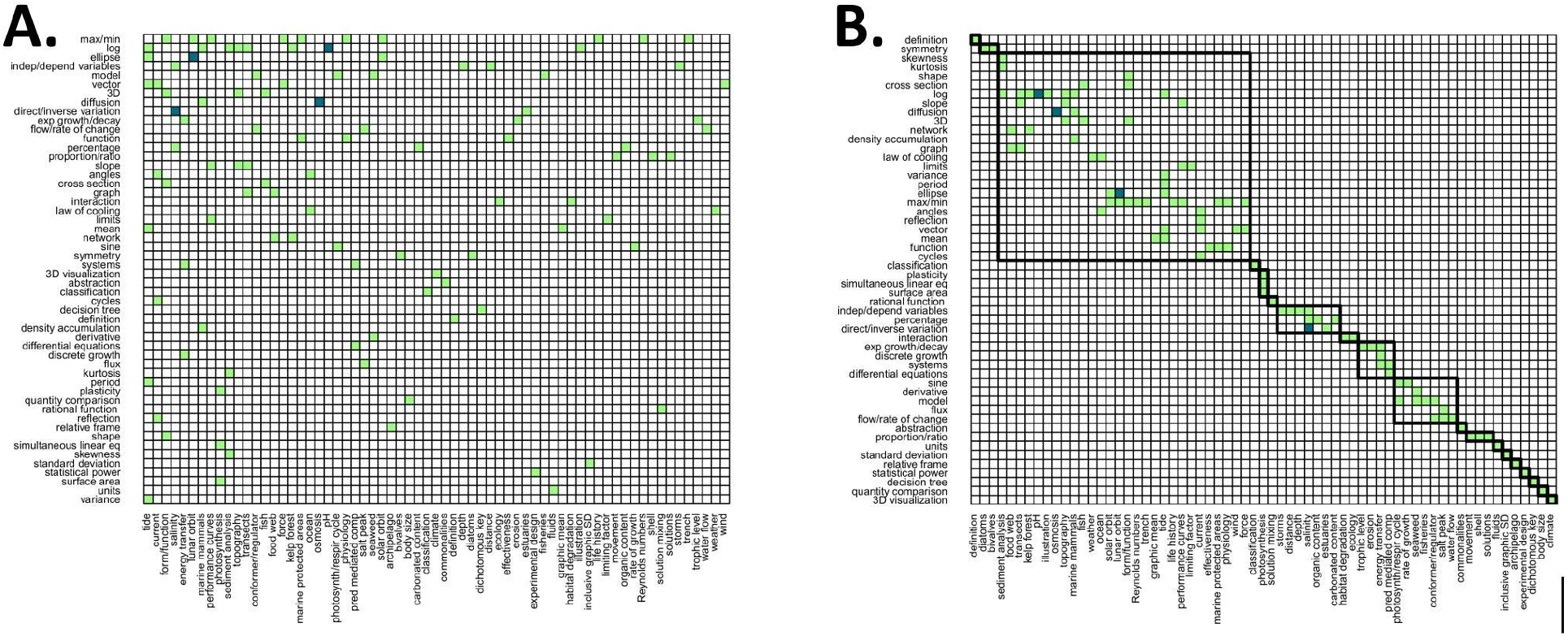
Grid plots of the math-marine concept network produced via ‘visweb’. White squares indicate a co-occurrence count of 0, light green indicates 1, and dark teal indicates 2, which was the highest observed. There are only four pairs with an edge size of two: ellipses and [planetary/lunar] orbits; salinity and direct/indirect variation; diffusion and osmosis; and log and pH. A is plotted with ‘type= “nested”‘ which sorts nodes by row and column sums, allowing for assessment of nestedness and concept connectedness. B is plotted with ‘type= “diagonal”‘ which plots the highest number of interactions along the diagonal and identifies compartments.

The compartment clustering approach in Figure 4B helps identify possible directions for an interdisciplinary module of mathematical and marine biology concepts. For example, we might create a module combining dependent/independent variables, percentages, and direct/indirect variation math concepts with solution mixing, distance, depth, and salinity. Conceptualizing the collaborative education space as a mutualistic bipartite network leads us to possible emergent directions for interdisciplinary curriculum development.

Figure 4 also indicates a relatively sparse interaction matrix. Connectance is the number of connections observed out of the number possible (Jordano, 1987) and in our case is 3.14%. While we do not know the connectance between mathematics and biology as fields, typical plant-pollinator networks average 30% connectance (Jordano, 1987). We know our sampling method falls short of seeing all connections. For example, for example it may change simply by including the laboratory component in the analysis or include multiple courses in the analysis.

Even if we are not sure what a target connectance value should be, we could use it as a metric for measuring and tracking our success in reaching interdisciplinary teaching. One caveat is that there is likely an optimal connectance value for any mutualistic network - more edges does not necessarily mean more network stability, and larger networks have smaller connectance values (Jordano, 1987). We conceptualize network stability here as a state of maximum entropy (Meysman & Bruers, 2010) or minimal uncertainty of information. In the context of interdisciplinary relationships, this may describe the state of links between disciplinary ideas or topics for which minimal information can determine the system (Çengel, 2021). Essentially, the disciplines themselves, or your mental organization of them is as efficient as possible in remembering the minimal number of connections needed to perform maximally in the future.

These ideas align with knowledge systems theory (diSessa & Sherin, 1998; Sabella & Reddish, 2007), but would be stronger if we could observe the network decreasing in connectance over time, for example over subsequent years of class observation.

We can also look for disciplinary ways to support interdisciplinary teaching. In Figure 5, we introduce a competition plot, which examines nestedness with more of a concept-focus rather than a webwide-focus. This makes it useful for considering novel ways of approaching the ‘lower’ level with the specific goal of enhancing understanding of the ‘upper’ level. The network graph in Figure 1 indicates that maximums/minimums and log are vital math concepts for marine biology. However, the competition plot tells us that if we want to teach about maximum and minimums in a holistic manner with the goal of supporting and integrating with marine biology, we could support the teaching of ellipses and vectors. Ellipses, surprisingly, are not part of all precalculus courses and vectors are largely covered in the physics curriculum, so this could direct additional disciplinary conversations within mathematics. In contrast to maximum and minimums, proportions and ratios is another relatively large node, but it is entirely self-contained. It has no clustering to other math concepts through a common marine concept.

**Figure 5.**
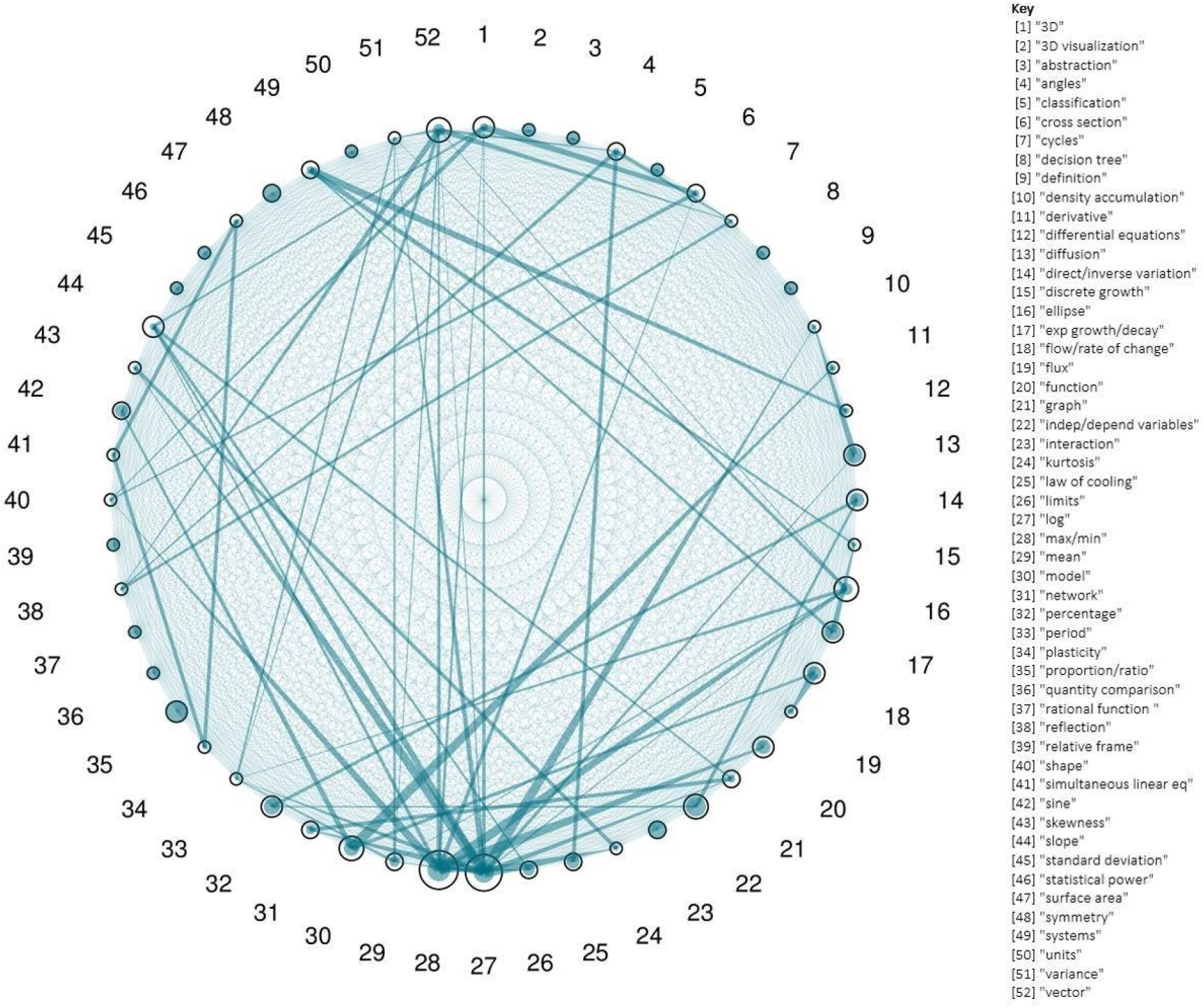
Competition plot with key of math (‘lower’ level) concepts produced via ‘plotPAC’. Black circles are nodes. The amount of teal fill in each circle indicates level of ‘specialization’. Teal lines indicate connections between nodes through marine (‘higher’ level) concepts.

## CONCLUSION

Many of the limitations of this methodology are the same as limitations for other observing ecological bipartite networks. The way we sampled our interaction network matters (Gibson et al., 2011). A more complete web of interactions would be had, for example, if we observed every marine biology class in the curriculum or across institutions. In addition, the time at which you inventory a network matters. We recommend these as potential future investigations. However, the context-specific nature can be viewed also as a strength of this approach. This framework understands interdisciplinary teaching practice as complex and adaptive and facilitates new approaches to explore how instructor practice might change over time as a result of the instructors, interventions, conversations, and other experiences.

A limitation potentially more unique to the interdisciplinary context is the ability to code the qualitative data. Recognizing when a mathematics reference has occurred in the biology classroom may be more nuanced than identifying when a plant has been visited by a pollinator. It may take some familiarity with the language across disciplines (Redish & Kou, 2015). We recommend further studies that explore the interrater reliability of this coding, particularly as it relates to disciplinary training and grounding.

Despite the limitations, our exploration of marine biology education and mathematics education as a mutualistic coevolutionary system yielded productive insights about supporting interdisciplinary teaching. Teaching practice experience may be affected by different individual interventional strategies. We found two key ways that significant math was infused into this class: 1) prescribed: use of guest lectures/pre-developed resources and 2) emergent: through marine topics that were close to the instructor’s area of research, training, and/or comfortability as defined by Henderson et al. (2011). These results seemed intuitive to the marine biology instructor. Form and function were an emphasis in their dissertation research area, and they intentionally choose a high-quality video to help explain concepts behind tidal and orbital physics.

The data collection and in-class collaboration process itself led to new curricular conversations and collaborations on the part of both math and marine biology instructors. Diaz Eaton’s participation in the SUMMIT-P project gave her better insight into authentic and meaningful marine biology motivations for calculus-related math ideas. For example, invasive green crabs are a concern in local coastal waters, so it provides a meaningful context to explore carapace length, total length, and age data for native and invasive crabs to explore differential equation growth models. The marine biology instructor was surprised to learn that vectors were not discussed in mathematics courses, but in physics, and we discussed whether physics should be required - or if not, teaching vectors more intentionally.

In addition to the sheer number of mathematics topics in the introductory marine biology course studied, we also found that there are significant challenges marine biology instructors and students may face inherently associated with the structure of mutualistic networks i.e., heterogeneity. The variety of mathematics concepts making one-time appearances in a course may make it difficult for marine biology instructors to fully prepare students for all mathematical skills necessary. Diaz Eaton was initially surprised about the emphasis of optimization and the appearance of conic sections (i.e., ellipses), the latter of which had been cut from the precalculus curriculum.

Theorizing partner disciplines as mutualistic and coevolving also helped identify ways to capture emergent points for potential new interventions. The nested grid plots (Figure 4B) were extremely useful to identify new connections across courses: ellipses could be used as an optimization example in the context of lunar orbits, or a project could be to model salinity levels as a function of water depth. Unfortunately, the primary investigator moved institutions and was unable to develop these ideas further. However, both the collaborative process as well as the network approach each provided useful directions to move forward interdisciplinary curricular collaborations. Finally, network indices, like connectance, may be helpful in evaluating the development of interdisciplinary teaching practice or instructor learning organization over time between course networks. These opportunities fully embrace the complex adaptive nature of the mutualistic system between mathematics and biology.

## ACKNOWLEDGEMENTS

This study is supported by a subaward through NSF IUSE #1625771 and #1822451, Bates College, and the William and Flora Hewlett Foundation. IRB approval UCIRB 2017-02. We thank the work of and conversations with subaward co-investigators, Dr. Perry and Dr. Wade, Unity College faculty and students, and the full SUMMIT-P collaboration team.

